# Statistical Pitfalls in Brain Age Analyses

**DOI:** 10.1101/2020.06.21.163741

**Authors:** Ellyn R. Butler, Andrew Chen, Rabie Ramadan, Trang T. Le, Kosha Ruparel, Tyler M. Moore, Theodore D. Satterthwaite, Fengqing Zhang, Haochang Shou, Ruben C. Gur, Thomas E. Nichols, Russell T. Shinohara

**Author notes:** All code can be found in https://github.com/PennBBL/brainAgeGapMistake. Denotes co-senior authorship (Thomas E. Nichols, Russell T. Shinohara). Denotes corresponding author, E-mail address, Postal address: Blockley Hall, 2nd Floor, University of Pennsylvania, 423 Guardian Drive, Philadelphia, PA 19104-6021.

## Abstract

Over the past decade, there has been an abundance of research on the difference between age and age predicted using brain features, which is commonly referred to as the “brain age gap”. Researchers have identified that the brain age gap, as a linear transformation of an out-of-sample residual, is dependent on age. As such, any group differences on the brain age gap could simply be due to group differences on age. To mitigate the brain age gap’s dependence on age, it has been proposed that age be regressed out of the brain age gap. If this modified brain age gap (MBAG) is treated as a corrected deviation from age, model accuracy statistics such as *R*^2^ will be artificially inflated. Given the limitations of proposed brain age analyses, further theoretical work is warranted to determine the best way to quantify deviation from normality.

**Highlights:**

- The brain age gap is an out-of-sample residual, and as such varies as a function of age.
- A recently proposed modification of the brain age gap, designed to mitigate the dependence on age, results in inflated model accuracy statistics if used incorrectly.
- Given these limitations, we suggest that new methods should be developed to quantify deviation from normal developmental and aging trajectories.

## 1 Introduction

In the past decade, there has been an explosion of research devoted to estimating individuals’ ages using features derived from magnetic resonance images (MRIs) of the brain (Franke & Gaser, 2019). From studies using diffusion-weighted features to complex functional connectivity metrics, the literature is extensive (Cole, 2020; Erus et al., 2015; Irimia, Torgerson, Goh, & Van Horn, 2015; Li, Satterthwaite, & Fan, 2018; Lin et al., 2016). While age is easily measured through more conventional means, assessing the appearance of the brain with respect to the natural patterns of development and aging provides a framework for dimension reduction; from hundreds of thousands to millions of MRI measurements, these models aim to provide the age of the brain for each subject as a convenient summary measure. The predicted age from these models has been coined “brain age”, and the difference between age (sometimes referred to as “chronological age”) and brain age is typically referred to as the “brain age gap”. Predicted ages are calculated using the following fitted model:

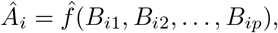

where *Á_i_* is the predicted age of the *i*th subject, *B_ij_* is the *j*th brain feature for the ith subject, and *f*(·) is some function of the brain features.

Brain age gap analysis was developed to address two major challenges in neuroscience and medicine: high-dimensionality, and individual risk assessment. Neuroimaging data are high dimensional, with the average T1-weighted scan containing approximately 1,200,000 voxels of brain tissue (Cosgrove, Mazure, & Staley, 2007). Importantly, different parts of the brain follow a variety of trajectories across the lifespan (Coupé, Catheline, Lanuza, & Manjón, 2017; Gennatas et al., 2017; Kennedy et al., 2015). Therefore, in order to better predict age, it is beneficial to use more brain features that complement each other (Varikuti et al., 2018). The main motivation, however, behind brain age gap analyses has been to develop a single number to represent an individual’s deviation from some normal trajectory (de Lange & Cole, 2020). This is an admirable goal, since deviating from a normal trajectory may be indicative of or predictive of debilitating disorders (Marquand, Rezek, Buitelaar, & Beckmann, 2016).

Researchers often test whether members of a group tend to have their age overestimated compared to a control group, striving to assess whether the disorder is associated with the brain aging prematurely or lagging behind. For instance, Chung et al. (2018) asked if those at clinical high risk for psychosis had a larger brain age gap than healthy controls, and Liem et al. (2017) asked if the brain age gap differed across groups with varying degrees of objective cognitive impairment. Typically, these models are developed using regression or machine learning in one dataset, and are evaluated in a test set. The cross-validation process involves dividing the training set into *k* folds, estimating the model parameters on *k* – 1 folds, applying the fitted model to the remaining fold, and repeating until every participant in the training set has a predicted age. This procedure helps avoid over-fitting and reporting an inflated model accuracy statistic. Finally, the trained model is applied on a separate test set to predict age of each individual based on their brain features.

In this article, we note that the brain age gap, and a recently proposed modified version of it (Beheshti, Nugent, Potvin, & Duchesne, 2019; Chung et al., 2018; Liang, Zhang, & Niu, 2019; Smith, Vidaurre, Alfaro-Almagro, Nichols, & Miller, 2019), are not up to the task of quantifying deviation from a normal trajectory. The brain age gap is a linear transformation of an out-of-sample residual (subsequently referred to as a “prediction error”). As such, it is dependent on the outcome variable (i.e., age) (Le et al., 2018). Therefore, differences in the brain age gap between groups may be due to differences in the brain, or due to differences in the age distributions across groups (Le et al., 2018; Smith et al., 2019). A recently proposed solution to this problem — regressing the brain age gap on age and taking the residuals from this model as a modified brain age gap that is orthogonal to age — creates new problems. In particular, if this new prediction error is treated as a deviation from a subject’s age, which it is not, metrics of model accuracy will be severely inflated.

## 2 Known Limitations of the Brain Age Gap

Brain age gap analyses have historically been based on the assumption that the difference between age and predicted age does not vary as a function of age; however, recently several groups have pointed out that this assumption is false (Le et al., 2018; Liang et al., 2019; Smith et al., 2019). Smith et al. (2019) pointed out an extreme case of this error: when age has truly no relationship with brain features, the difference between age and predicted age (“brain age gap”) is a linear function of age, which implies that age explains 100% of the variance in the brain age gap. Smith et al. (2019) note that any subsequent analyses studying the relationship between this gap and other metrics is equivalent to relating a linear transformation of age to other metrics.

To flesh out the gravity of this observation, consider an example: If age does not vary as a function of any of the brain parameters, all coefficients, aside from the intercept, will be close to zero with high probability, and the intercept will be close to the mean age of the training sample. Let *A_i_* be the age of the *i*th subject, *B_ij_* the *j*th brain feature for the *i*th subject, *ϵ_i_* random error, and *Ā* the mean age of the training sample. The brain age model is thus:

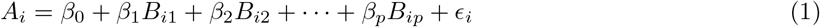

And the fitted values are:

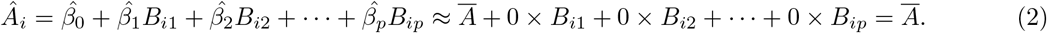

For simplicity, let’s assume that the coefficients are estimated to be exactly zero. Suppose the mean age of the training sample is 10 years old. Every person will have an estimated age of 10, so their brain age gap, *Â_i_* – *A_i_*, will be 10 – *A_i_*. Thus, the brain age gaps are as follows: 15-year-olds have a brain age gap of −5, 10-year-olds have a brain age gap of 0, 5-year-olds have a brain age gap of 5, etc. Older participants are estimated as being younger than they are, and younger participants as older. The brain age gap is a linear transformation of a residual (i.e., 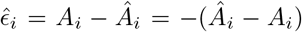), which by definition varies as a function of the outcome variable, in this case age. If the brain features are linearly independent of age, then testing for differences in the brain age gap is equivalent to testing, “Is the mean age of group A different from the mean age of group B?” When testing for differences on the brain age gap in general, the question being asked is similar to “Controlling for the brain features, is the mean age of group A different from the mean age of group B?” Because regression on the residuals of a previous model is not equivalent to multiple regression, this description is not quite correct (Chen, Hribar, & Melessa, 2018; Freckleton, 2002). Thus, interpretation of these residuals is difficult.

Even if age varies as a function of the brain parameters, the predicted age for every subject will still be shrunk towards the mean age of the training sample. This is referred to as regression towards the mean, and was first documented by Sir Francis Galton in 1886 (Bland & Altman, 1994). As Liang et al. (2019) noted, this phenomenon is a common feature of many good models. Therefore, older subjects will have negative brain age gap estimates on average simply because they are older, while younger subjects will have positive estimates on average.

It is important to note that regression towards the mean is not a failure, but a feature, of regression and related methods. If there is any randomness in a process, observations will tend towards the mean of the outcome variable rather than remain as extreme as they were upon initial sampling (Stigler, 1997). Regression towards the mean is a feature of regression that is actively useful for prediction. Since age is known with certainty, the notion of predicting it makes the construction of a residual awkward. Thus, as we continue to use age prediction as a means to reduce dimensionality, it is important to understand the limitations of using age as an outcome variable and subsequent associated analyses. Recognizing the dependence of the brain age gap on age, researchers have begun to develop methods to mitigate the age-dependence of the brain age gap (Beheshti et al., 2019; Le et al., 2018; Smith et al., 2019). Unfortunately, a misuse of residuals persists, resulting in a systematic overestimation of model accuracy.

## 3 Risks of Using a Modified Brain Age Gap

To mitigate the residuals’ dependence on age, some researchers apply the following algorithm (Beheshti et al., 2019; Chung et al., 2018; Liang et al., 2019; Smith et al., 2019) (see the appendix for details on Beheshti et al. (2019)’s method). First, a training sample is used to estimate a mapping *f*(·) from brain features to age. Then, for a left out subject *i* with brain data *B*_*i*1_, *B*_*i2*_,…, *B_ip_*, the predicted age (“brain age”) is estimated as *Â_i_*:

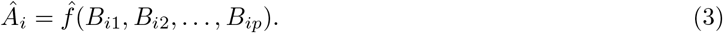

Then the *i*th subject’s brain age gap (BAG) is

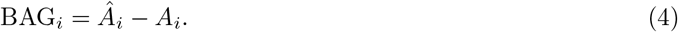

Recognizing the brain age gap’s dependence on age, the researcher poses a linear model of the brain age gap on age:

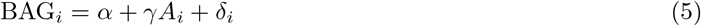

where estimated parameters 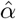 and 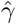 are found from a regression using training data, and *δ_i_* is random error. Thus, the effect of age is removed, producing the modified brain age gap (MBAG):

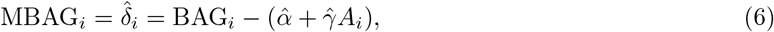

which, as the prediction error from model (5), is approximately uncorrelated with age (only exactly uncorrelated if test data is used to estimate *α* and *γ*). Because MBAG has been interpreted as a corrected residual, MBAG is added to (or subtracted from; equivalent in correlation, see Supplement) age. This new variable is then referred to as the corrected predicted age:

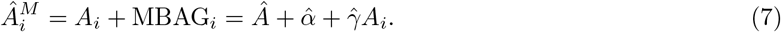

Because the researcher perceives this predicted age as corrected, they correlate it with age to assess their model’s accuracy in predicting age. We will refer to 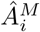 as the “modified predicted age” and will show below why this age estimation is flawed.

MBAG is by no means a more accurate measure of an out-of-sample residual, or prediction error (i.e., the “brain age gap”). The brain age gap itself is *more* dependent on age the *less* the brain features are associated with age. Again, consider the extreme case where age is independent of the brain features. Then, the brain age gap is *completely determined* by age, as explained in the previous section. If MBAG is treated as an estimate of the deviation from age, the reported model accuracy (e.g., 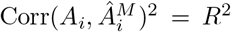) will always be inflated relative to the true model accuracy, and often drastically so (see Table 1 for details on papers that have reported inflated model accuracy statistics). When age has no true dependence on the brain features, the population covariance between age and predicted age, *Â_i_*, is zero. But when MBAG is treated as the deviation from age, *A_i_* + MBAG_*i*_, age and modified predicted age, 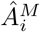, have an approximately *perfect correlation of 1*.

**Table 1:** Papers reporting inflated model accuracy statistics.

| Paper | Before Modification | After Modification |
| --- | --- | --- |
| Beheshti et al. (2019) | $\text{Corr}(A, \hat{A})^2 = .38$ | $\text{Corr}(A, \hat{A}^M)^2 = .88$ |
| Chung et al. (2018) | $\text{Corr}(A, \hat{A})^2 = .66$ | $\text{Corr}(A, \hat{A}^M)^2 = .84$ |
| Liang et al. (2019) | $\text{MAE} = 1.57$ | $\text{MAE} = 1.32$ |
| Smith et al. (2019) | $\text{Corr}(A, \hat{A}) = .06$ | $\text{Corr}(A, \hat{A}^M) = .99$ |

Note: MAE = Mean Absolute Error.

In fact, the inflated correlation can be directly computed as a function of the sample estimates of the covariance between age and predicted age, the variance of age, and the variance of predicted age (see Supplement for derivations):

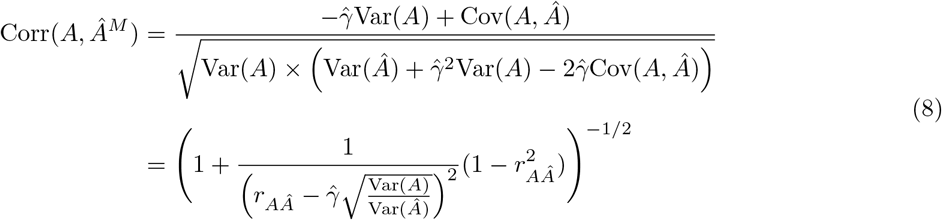

If 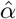 and 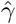 are estimated in the test set, equation (8) can be further simplified:

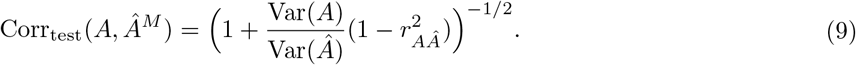

The equation can be simplified even further if *Â* is a linear estimator:

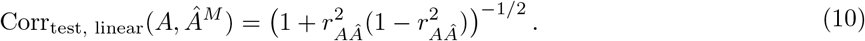

To illustrate the inflated correlation effect and confirm that equation 8 is correct, a series of simulations were run to compare the transformations that researchers describe performing to the above equation using R version 3.6.2 (R Core Team, 2019). Training and testing sets of 10,000 samples were simulated from each of a series of bivariate normal distributions, where the true correlation between age and brain was varied between 0 and 1, with the correlation between age and the modified predicted age, 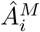, in the test set being the key outcome measure recorded. All model parameters were estimated in the training set. Since there is only one brain feature, the correlation between age and predicted age is the same as the correlation between age and brain. Results using a single brain feature are detailed in Figure 1. A single brain feature was used so as to have easy control over the correlation between age and predicted age, but note that this result generalizes to any number of brain features. For a set of correlations between 0 and 1, the correlation between age and the modified predicted age, 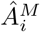, was calculated using the theoretical formulation in (8) (black line), and the inflated correlation was obtained using the previously described transformations (pink dots). The identity line is displayed to aid in visualizing that the inflated correlation is larger than the true correlation. The simulations confirmed that the theoretical formulation in (8) is equivalent to the transformations researchers have described. In addition, Figure 1 illustrates that the degree of inflation is much greater for models that have lower values of Corr(*A, Â*) than for models that have higher values of Corr(*A, Â*).

**Figure 1:**
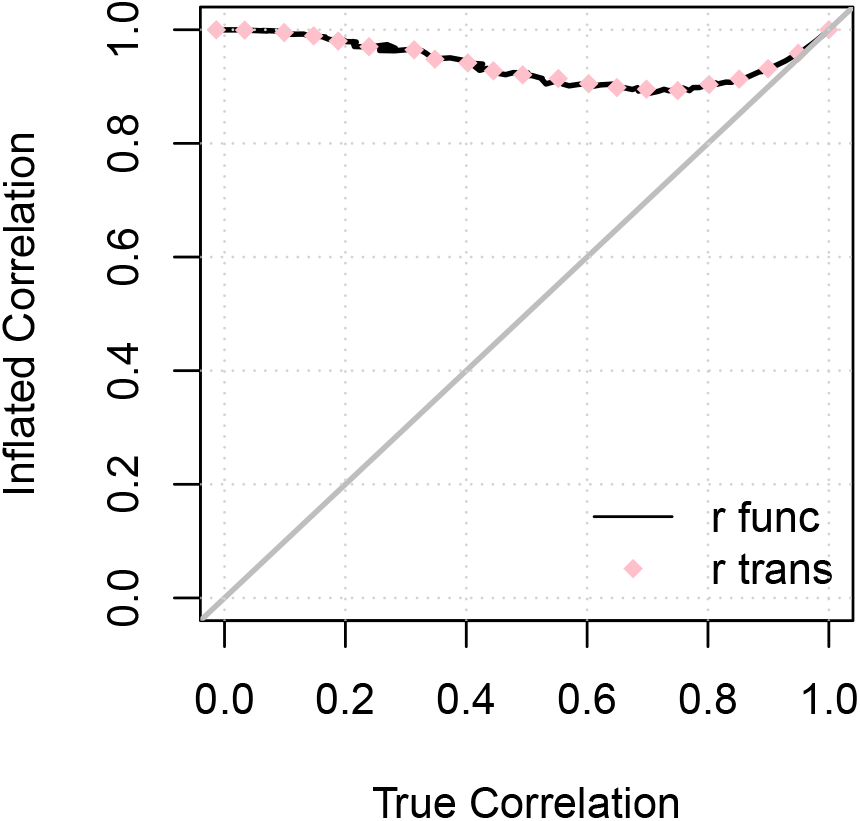
Inflated correlation, Corr(*A, Â^M^*), is a function of the true correlation, Corr(*A, Â*). The inflated correlation is the correlation between age and the modified predicted age. The true correlation is the correlation between age and predicted age. To illustrate that the series of transformations that researchers perform is equivalent to (8), correlations using both are plotted. *r*_func_ is using (8), and *r*_trans_ is using the series of transformations. The identity line is displayed.

Additional analyses were run using the Philadelphia Neurodevelopmental Cohort (PNC) to illustrate the findings in brain MRI data. Sample details, neuroimaging protocols, and processing can be found in Calkins et al. (2015), Gur et al. (2020), and Satterthwaite et al. (2014). Briefly, participants ages 8-22 were recruited through their primary care providers in the Philadelphia area. Subjects were excluded for the purposes of these analyses if their cognitive assessment was conducted more than a year before or after their neuroimaging data was collected, or if their structural image did not pass stringent quality assurance measures. 132 regional volume values were extracted using the Advanced Normalization Tools software package (Tustison et al., 2013; Wang & Yushkevich, 2013).

Elastic net models to predict age were built on youths ages 8-22 without a history of mental illness (“typically developing”). Hyperparameters were chosen using repeated five-fold cross validation on the typically developing youth as implemented in the ‘caret’ package, version 6.0-86 (Kuhn, 2012). Then, a linear regression of BAG on age was fit in the typically developing subjects (N = 317). Using the fitted values for the parameters from these models, the transformations previously described were applied to youth who met screening criteria for lifetime instance of a mental illness (N = 862). This real data example confirmed the theoretical and simulation findings (see Figure 3). Prior to any modification, the correlation between age (*A*) and predicted age (*Â*) was .773. After applying the modifications, the correlation became .884. There were no differences between the typically developing youth and youth with a history of mental illness on age (*t* = –1.05, *p* = 0.29), the brain age gap (*t* = 0.72, *p* = 0.47), or MBAG (*t* = 0.09, *p* = 0390). Age and performance on the complex cognition tasks were highly associated (*r* = 0.54, *p* < .0001). After regressing the brain features out of age and multiplying by negative one - or constructing the brain age gap - this association weakened (*r* = –0.30, *p* < .0001). MBAG and performance on the complex cognition tasks were not associated (*r* = –.01, *p* = 0.71). These results indicate that the association between cognition and the brain age gap are driven by the association between age and cognitive performance. Prior work highlighting group differences and correlations between brain age metrics and other variables should be examined in light of these results.

## 4 Conclusion

We have shown that predicted age estimates (“brain age”) based on a regression adjustment of the brain age gap result in a correlation between a modified predicted age and age never falling much below 0.9 regardless of the original predicted age and age correlation. Further, the interpretability of MBAG itself is limited by the fact that it is a prediction error from a regression to remove the effects of age from a residual obtained through a regression to predict age. By virtue of these limitations, we suggest that the brain age gap and the modified version may not provide useful information about precocity or delay in brain development. In light of this, we suggest that methods be developed specifically to answer questions about similarity between brains of different age groups and diseased states.

Many other transformations have been developed to mitigate the downstream effects of BAG’s dependence on age (de Lange & Cole, 2020). Some are not susceptible to the inflated correlation issue described in this work. Methods include scaling the predicted age by the slope and intercept from the regression of predicted age on age (see (5) in de Lange and Cole (2020)), and including age as a covariate when testing for group differences in BAG (Le et al., 2018). The former results in a new BAG estimate that is uncorrelated with age, and the latter ensures that any group differences found on BAG will be linearly independent of age. Note that, if all models had been built on the test set, controlling for age when testing for group differences on BAG is the two-step regression equivalent of including age as a covariate in a multiple regression with brain features predicting age. The real question then becomes: to what extent do these methods quantify advanced or delayed brain development? This question warrants further theoretical investigation.

**Figure 2:**
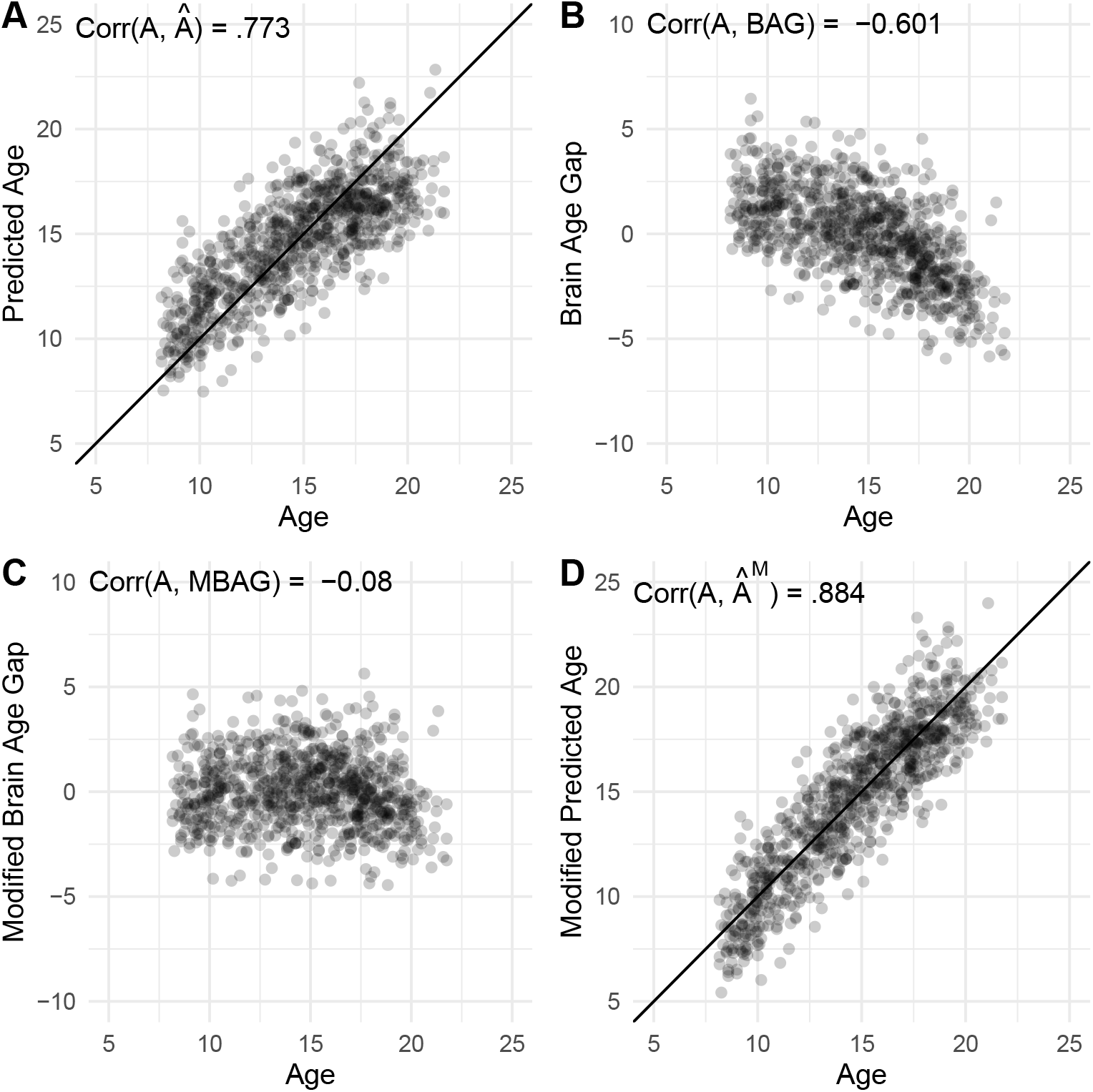
The inflated correlation finding was replicated in the Philadelphia Neurodevelopmental Cohort. Plotted are values for age, predicted age, brain age gap, modified brain age gap and modified predicted age in the subset of participants who met screening criteria for an instance of mental illness in their lifetime. The identity line is displayed in panels A and D.

Future research should also determine appropriate analytic methods to answer whether the brains of patients with disorders are more similar to older healthy controls’ than age-matched healthy controls’ brains, and to evaluating the extent to which analyses of residuals as deviations from some trajectory exist in the literature. Thus far, we are aware of a similar trend of predicting age using genetic features in attempt to document differences in precocious and delayed genetic development (Sumner, Colich, Uddin, Armstrong, & McLaughlin, 2019; Wolf et al., 2018). In the meantime, while previous studies have suggested that the brain age gap be used as biomarker in clinical trials (Cole et al., 2018), our findings suggest that further methodological work is warranted.

## 5 Acknowledgements

E.R.B. conceptualized the work, wrote derivations, and wrote the manuscript. A.C., T.E.N., and R.T.S. wrote derivations, and provided feedback on the framing of the work. R.R., T.T.L., F.Z. and H.S. provided statistical insights and edited the manuscript. T.D.S. oversaw the processing of the neuroimaging data and edited the manuscript. K.R., T.M.M., and R.C.G. provided inspiration for, and assisted in the framing of the work. We thank Dr. Michael Stein, members of the Brain Behavior Laboratory and attendees of Data Club for their questions and feedback.

## Beheshti et al. (2019) correlation

Beheshti et al. (2019) suggest subtracting 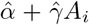 from *Â_i_*, and calling this new value the corrected predicted age:

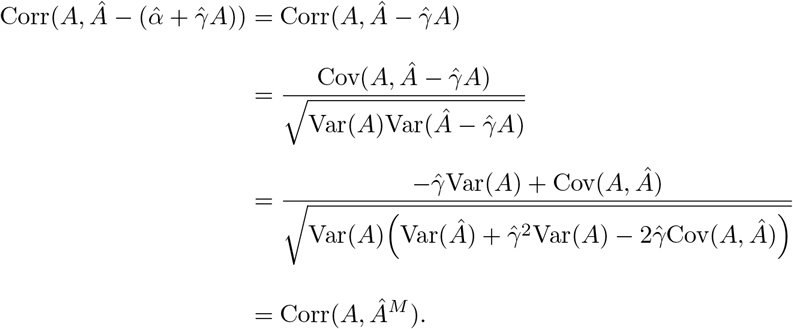

Therefore, their method is equivalent to Eqn. 8.

## Adding and subtracting MBAG from age results in the same inflated correlation with age

Moditified predicted age has been calculated in the literature as either MBAG_*i*_ = *A_i_* −BA*G_i_* or MBAG_*i*_ = *A_i_* + BAG*_i_*. In both cases, the main results from the paper applies since the correlation between age and the modified predicted age using either formula is the same. We have that

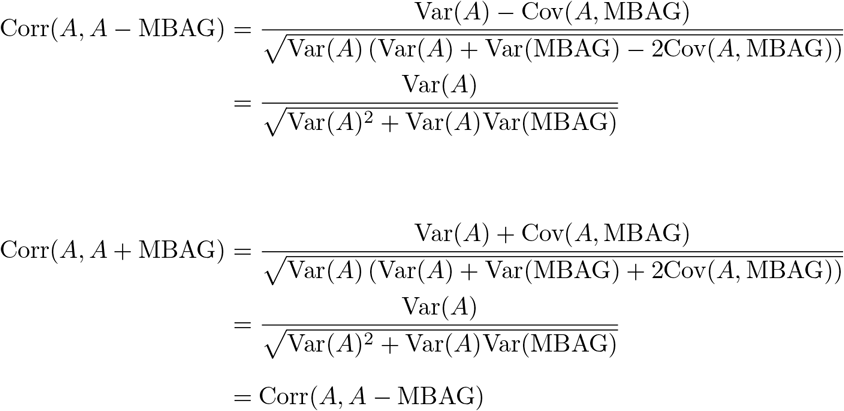

which follows from the fact that MBAG_*i*_ is a residual from regression of BAG*_i_* on *A_i_* and thus MBAG_*i*_ is orthogonal to *A_i_* or equivalently, Cov(*A*, MBAG) = 0. Note that this result is only approximate when the regression of BAG on age is done in the training set.

## Derivation of Equation (8)

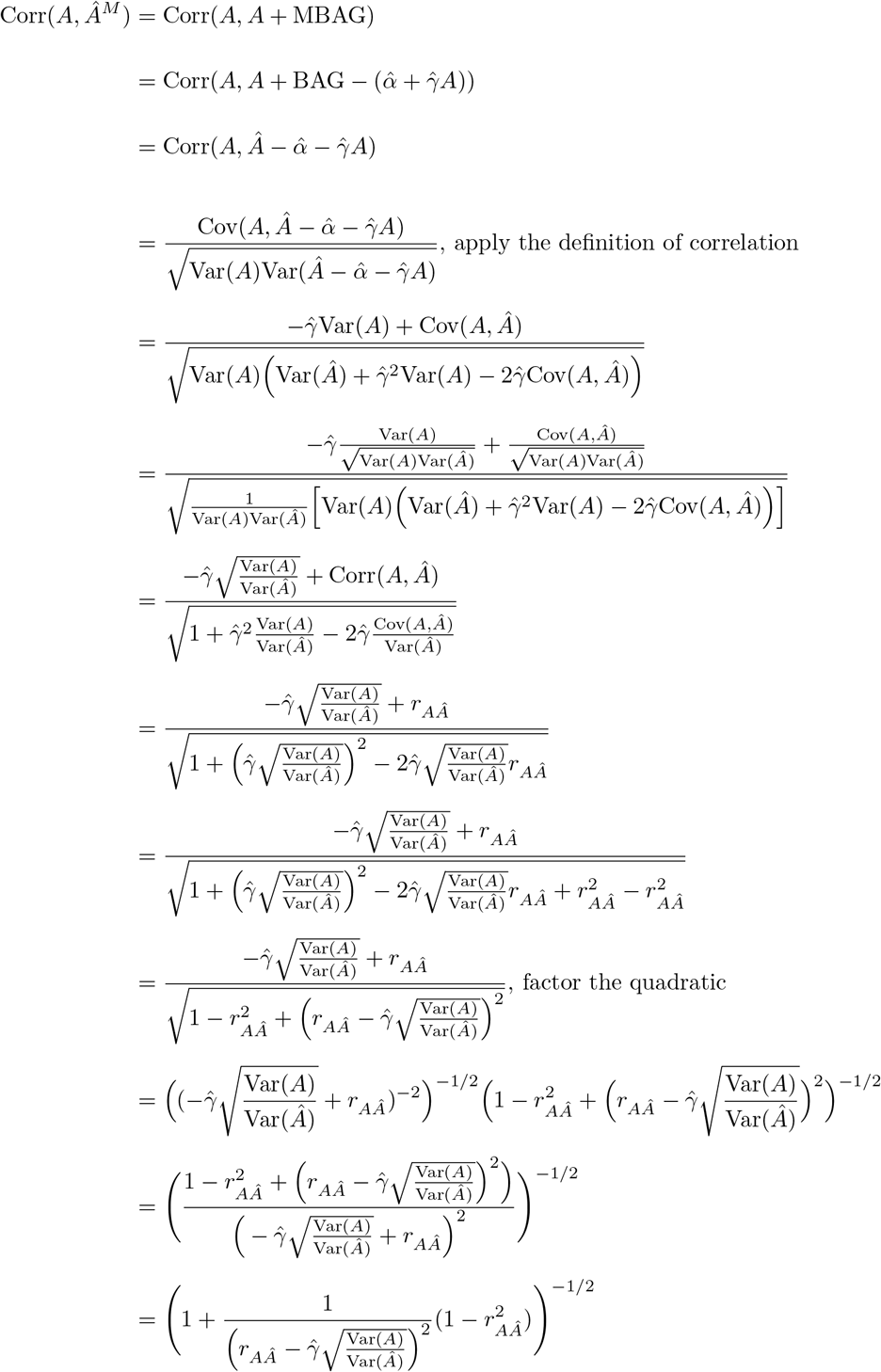

## Derivations of Equations (9) and (10)

The following derivation involves the algebraic manipulation of the sample estimates and not expectations. Assuming that the linear regression of BAG on age has been estimated with the testing data, then

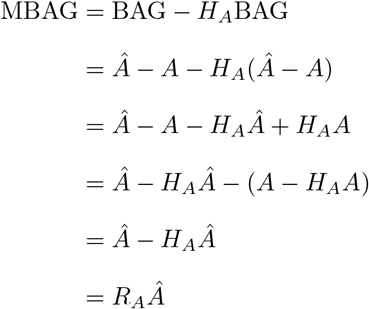

where *H_A_* = **A**(*A*^*T*^ **A**)^−1^**A**^*T*^ is the hat matrix for the regression on age, **A** = [**1** *A*], and *R_A_* = *I* – *H_A_* is the corresponding residual forming matrix.

We first note that

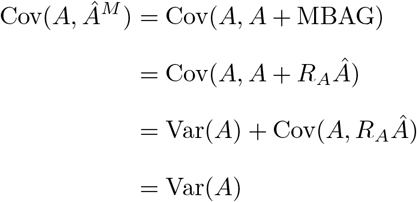

where the last equality holds due to the orthogonality of *A* and *R_A_*.

Then, consider:

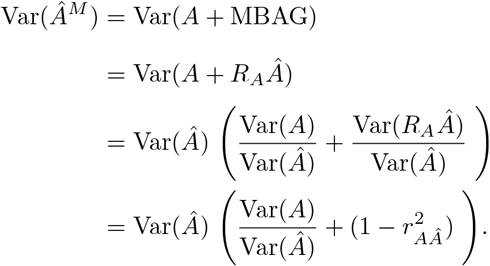

Then, Eqn. (9) is found as

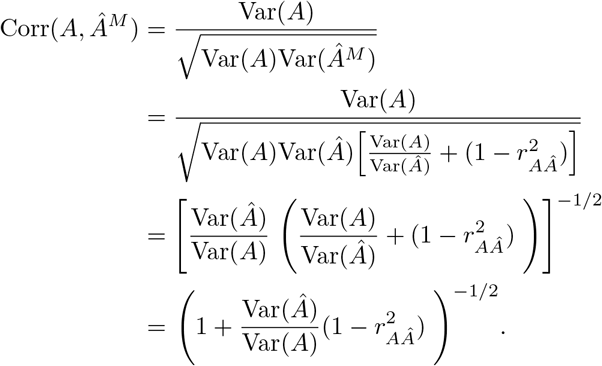

For insight on the Var(*Â*)/Var(*A*) term, note that shrinkage will generally mean this term is less than one. Moreover, if *Â* were found with a linear regression on the testing data, i.e. *Â* = *X*(*X^T^X*)^-1^*X^T^A* where X are brain features, then this ratio is exactly the squared correlation,

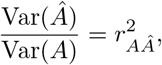

producing Eqn. (10).

In this setting, when both brain age and MBAG are determined from testing data using linear regression, the correlation of *A* and *Â^M^* can never fall below 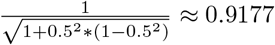. Of course, in practice, held-out training data is used to learn the brain-age relationship, so a regression prediction would instead have the form 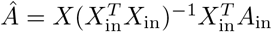, where *X*_in_ and *A*_in_ are held-in training data, but 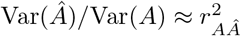 still provides a useful starting point for exploring the parameters in the expression for Corr(*A, Â^M^*).

Finally, note that the equality of Var(*Â*)/Var(*A*) and 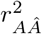 holds not just for linear regression, but any linear estimator. Specifically, if there exists an idempotent 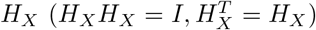 such that **1***^T^H_X_* = **1** and *Â* = *H_X_A*, then

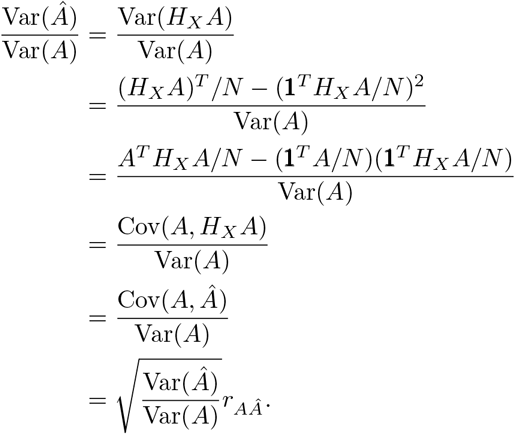

And thus

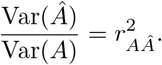

